# Prevalence and Associated Factors of Agricultural Technology Adoption and Teff Productivity in Basso Liben District, East Gojjame Zone, North west Ethiopia

**DOI:** 10.1101/2020.10.28.358770

**Authors:** Demelash Abewa Elemineh, Hayimero Edmealem Merie, Mulusew Kassa

**Author notes:** **correspondence author:** Demelash Abewa Elemineh, Department of Statistics, Debre Markos University, Debre Markos, Ethiopia.

## Abstract

Teff productivity and Teff technology adoption in Ethiopia is low and it requiring immediate attention for policymakers and decision-makers. This study was conducted to identify the determinant factors that affect Teff technology adoption and Teff productivity in Basso Liben district, East Gojjam Zone, Northwest Ethiopia. A cross-sectional study design was conducted among 190 households. Multivariable linear and logistic regressions were employed to identify the factors associated with Teff production and Teff technology adoption respectively. Of a total of 190 households considered in the study, 77.9% were Teff technology adopter. Household head gender (male) (OR=7.644), family size (OR=1.149), age of household head (OR=0.873), row planting (use)(OR=257.2), credit access (yes)(OR=3.141), manure(use)(OR=0.042) were significance associated with Teff technology adoption in the study area. Age of household head (*β* = 0.079), Education level (primary)(*β* = −0.612), total land holding (*β* = 5.107), annual income(*β*=0.0051), extension service (no)(*β* = −0.635), row planting (yes) (*β* = 1.409), organic fertilizer (no)(*β* = −0.946) were significance associated with teff production in the study area. In this study, a low prevalence of agricultural technology adoption and Teff production and various associated agricultural technology adoption and Teff production factors have been identified in the study area. Thus, the concerned stockholders should intervene in agricultural technology adoption and Teff production via different extension service and by considering household size, community-based household head education, and efficient use landholding in hectare.

## Background of the study

Teff is believed to have originated in Ethiopia & endemic to the country (1) and the crop occupies over 2.8 million hectares of the country (25-30% of the total cultivated area) (2). However, given the relatively low yields of Teff, the total national production of Teff (3.5 million ton) was lower than maize (6.1 million ton) and sorghum (3.9 million ton) (3). Often considered an “orphan crop” -one which hasn’t received the same kind of international attention from agronomists and commercial grower’s Teff has yet to fully benefit from modern farming technologies or techniques. These constraints have kept the crop’s potential yields low, while driving the price of the grain out of range for many Ethiopian families. Boosting yields and production of Teff, however, has the potential to significantly impact the livelihoods of millions of smallholder farmers along with the country’s economy as well.

Agricultural production could be enhanced either by increasing the area coverage or by increasing the productivity of agriculture (4). However, in the context of Ethiopia land is scarce and hence working on the extensive margin is very difficult. Hence, the feasible way to achieve this goal is through increasing productivity by investing in agricultural technologies or improving the level of technical efficiency (5).

Amhara region is the second largest Teff producer in the country next to Oromia region. The crop in Amhara region is produced by 228,502 smallholders and 426 large scale farmers on 184,648 ha. These farmers harvested about 5,159,33ton, with an average productivity of 2.29 ton/ha (6). Although the actual production of Teff is 2.29 ton/ha, the report of national Teff research commodity strategy 2016-2030, shows that the productivity of Teff can be increased by 4.34 ton/ha if farmers could adopt agricultural technologies (such as improved seed, row planting, herbicide and fertilizer). This means the production of Teff can be increased by 13 million tons in the country.

Agricultural technology’ includes all kinds of improved techniques and practices which could affect the growth of agricultural outputs (7). The most common agricultural technologies include improved varieties of seeds and farm management practices such as soil fertility management; weed and pest management and irrigation and water management. Adoption of these new technologies increases agricultural productivity, which can be seen through the outward movement of the production frontier, Such technologies are believed to be major factors for the success of the green revolution experienced by Asian countries (8).

The issue that requires an assessment is to what extent farmers are adopting agricultural technologies and what factors hinder small holders not to fully adopt the agricultural technologies in the way agricultural technologies were supposed to be delivered. Indeed there are few research conducted so far on this area in Ethiopia. Furthermore, to the best of our knowledge, Teff productivity and agricultural technology adoption is less clearly documented in the study area. Moreover, the statistical methodology used in the related Teff productivity and agricultural technology adoption literature was more qualitative and could not demonstrate the magnitude of the Teff productivity and agricultural technology adoption and could not explicitly specify the associated factors. Therefore, this study aims to evaluate the determinant factors that affect Teff technology adoption and Teff productivity in Basso Liben district, East Gojjam Zone, Northwest Ethiopia via binary logistic and multiple linear regression.

## Materials and Methods

### Description of the study areas

This study was conducted in Basso Liben district which is found in the northern highlands of Ethiopia stretching from 10°37 -10°38N latitude and 37°30-30°30 E longitude. The capital town of Basso Liben district is Yejube. It is located at a distance of 27 km from Debre Markos in south direction, 292 km from Bahir Dar and 317 km from Addis Ababa. The total area of the district is 113391.48 hectare. Its weather condition is 48% Woynadega, and about 54% kola with the latitude ranges 848-2417m above sea level. Most of kola part lies at Abay river gorge in which the area is owned by the Regional government of Amhara. The average rainfall is 900-1200mm per annual and the mean annual temperature is 15.5-20°c.

### Source and Study population

All farmers who live in Baso Liben district are the source population while all randomly selected farmers who live in the selected Kebeles would be the study population.

### Inclusion and exclusion criteria

All farmers, who live in the selected Kebele would be included. Those who were unconscious and mentally disabled would be excluded.

### Sample size determination

Sample size determination has its own scientific approach. But in this finding to determine the sample size, different factors such as research cost, time, and human resource, environmental condition, accessibility and availability of transport facilities were taken into consideration. By taking these factors consideration 190 household heads were selected.

### Sampling procedure

The districts were selected purposively based on Teff growing potential and improved Teff varieties have been introduced. In this study a two stage sampling technique was employed. The first stage was random selection of Teff growing Kebeles from the study area, followed by selection of sample households randomly. Hence, a total of 3 Kebeles (namely Yegelaw, Dendegeb and Enetemen) Teff growing Kebeles was randomly selected. Finally a total of 190 sampled households were randomly selected from the sampled Kebeles (Yegelaw=45, Dendegeb=85 and Enetemen=60 households)

### Dependent variable

Productivity of Teff: It is a continuous variable which represents the amount of Teff per hectare produced by household in the year 2018/19.

Teff Technology Adoption (yes, no): adopters are those households which use all Teff technologies such row planting, chemical, improved seed, extension service and fertilizer. Non adopters are those households which are not using the Teff technology comprehensively.

### Data collection methods

Primary data were collected using quantitative approach by means of household survey. The qualitative method of data collection was also employed. It consisted of in-depth open-ended interviews, direct observations and written documents. The interview method was mainly emphasized on group discussion and individual interviews were held to have reactions of the farmers relating to their detail experiences and their perceptions of Teff technology adoption and their priority problem why not they were adopted. Discussions with district experts of the agricultural office and key informants were also conducted. Before the administration of the structured and semi-structured interview schedules, exploratory farm surveys were conducted and the respondents were informed about the objectives of the survey. The interview schedules were pre-tested before the actual data was collected for clarity, acceptability, and flow among farmers who are out of the study area. Based on the findings from pre testing the questions that were difficult to answer were modified and amendments were made that the questions to make them fit to the context. Six enumerators and two supervisors were recruited. They were trained on the objective and contents of the interview schedule. The six enumerators conducted the interview in the local language, Amharic with the supervisor and researcher follow-up.

### Analysis/treatment of the data

The collected data was analyzed by using STAT version 13 Software and econometrics model for identifying factors that affect Teff Technology adoption logistic regression model was applied and for Teff productivity multiple linear regression model was also used.

### Econometric Model Specification

Multiple linear regressions are a statistical tool for the investigation of relationships between variables, usually to determine the causal effect of one variable upon another. Regression analysis estimates the conditional expectation of the dependent variable given the independent variables that is, the average value of the dependent variable when the independent variables are held fixed.

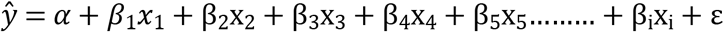

Where

*α*=Constant term,
β_i_= coefficient of multiple linear regression
εi = error term

Logistic regression analysis extends the techniques of multiple regression analysis to research situations in which the outcome variable is categorical. Then the conditional probability that the household is adopt teff technology given the X set of predictor variables is denoted by Prob (Yi =1|X) =Pi. The expression Pi has the form:

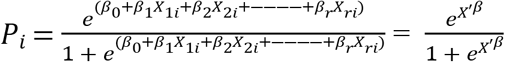

Pi = the probability of the i^th^ teff technology adoption
Yi = the observed survival status of household i
β is a vector of unknown coefficients.

## Result and Discussion

### Technology adoption characteristics of study participants

A total of 190 households from the Basso Liben districts, explicitly Yegelew, Dendegeb and Enetemen Kebeles, were considered in this study. The prevalence of Teff technology adoption varies by Kebele which was 71% for Yegelaw, 77% for Dendegeb and 85% for Enetemen. It shows that the proportion of Teff technology adoption was higher in Enetemen kebele, and to the contrary the proportion of Teff technology adoption was lower in Yegelaw kebele. Teff technology adopter and non-adopter were the two intervention areas in the households. The total proportion of households who use Teff technology adopter was 77.9% (Figure 1). This finding in line with previous reports from Basso Liben district 76.9% (9),. It was also relatively a low compared to previous reports from central high land of Ethiopia 79% (10). On the other hand the current result was higher than studies conducted in Ethiopia 54.44% (11), North east Ethiopia 17.8% (12), Central high land of Ethiopia 32.% (13), Amhara region 46.3% (14), South nation nationality people region 56.8% (15), Oromia region 63% and Malawi 41% (16). This difference, may be due to the fact that the variation of knowledge, attitudes and perceptions in relation to the benefits of the technology adoption play a key role in the decision to adopt (17).

**Figure 1:**
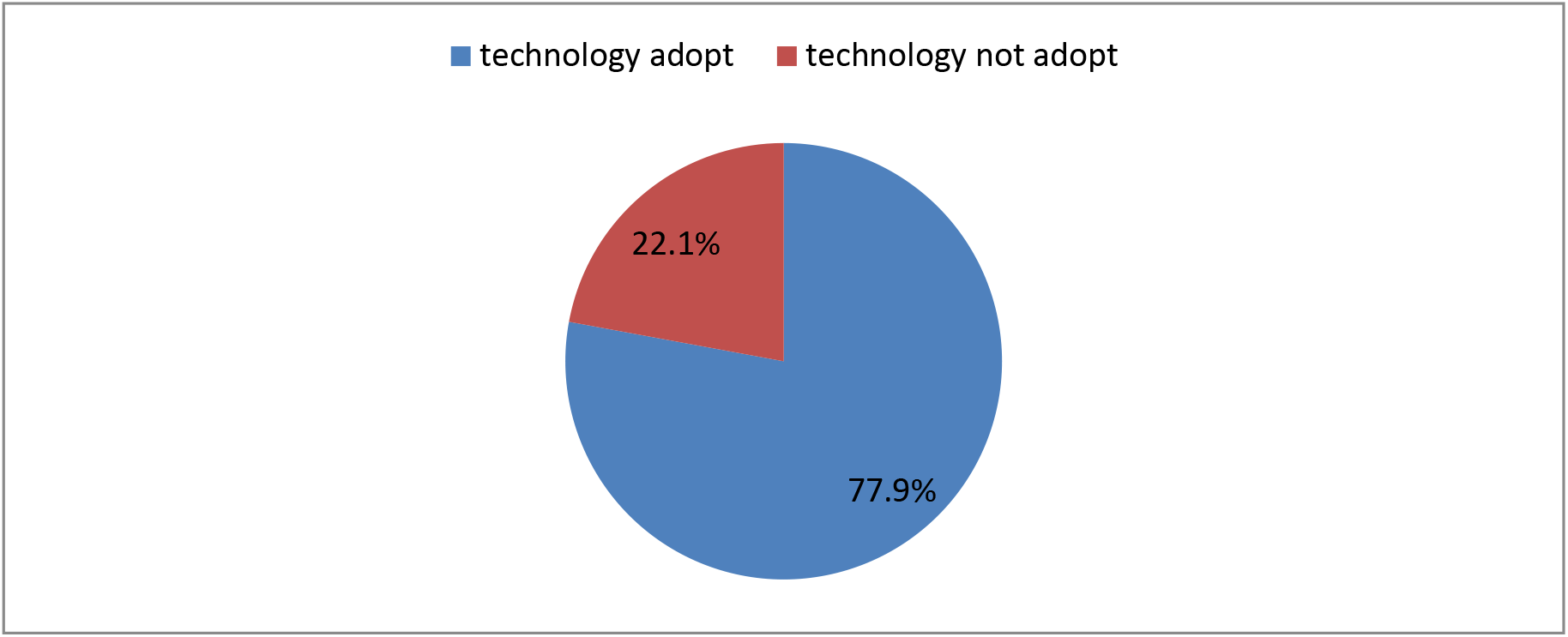
Teff production agricultural technology adopter and non adopter

### Socio demographic and economic characteristics of the farmer

In an agrarian society, household members are the major source of labor for agricultural activities (18). The average family size was 4.95 persons per household head with minimum 1 and maximum 8 persons per household. But there was a wide variation in family members among households. This finding is higher than the previous finding in Ethiopia 1.13 person per household (19) and in Ghana1.26 person per household (20). The average age of household head in the study area was 42.61 years old with minimum 28 and maximum 65 years. (Table 1). This finding is slightly lower than the previous report in Ethiopia 45.25 years old (19) and Western Ethiopia 45.07 years old (21). In the present study the total landholding of the sample households ranges from 0.00 to 4 hectare with an average figure of 1.08 hectare, which is slightly higher than the national average of 1.02 hectare (22) and lower than 1.17 hectare reported by Mesfin AH et al. (21). The average farm size in Ethiopia is less than two hectares reported by Sibhatu KT and Qaim M (23). The average annual off-farm income of the sample households were 4821.05 birr, which is higher than the national average annual off farm income 1162.70 birr (22).

**Table 1:** Socio-demographic and economic characteristics of the household head

| Variable | Mean | Std. deviation |
| --- | --- | --- |
| Age | 42.61 | 7.79 |
| Family size | 4.95 | 1.92 |
| Landholding(hector) | 1.78 | 1.08 |
| Amount of Teff production(quintal) | 24.23 | 13.66 |
| Annual off farm income ( birr) | 4821.05 | 7906.50 |
| Annual on farm income ( birr) | 48584.21 | 27018.52 |
| Amount of DAP (kilogram) | 72.44 | 50.33 |
| Amount of UREA (kilogram) | 47.07 | 44.10 |
| Amount of improved seed (kilogram) | 31.33 | 16.75 |
| Amount of herbicide(liter) | 0.70 | 0.74 |
| Amount of pesticide (liter) | 0.71 | 0.74 |

The average amount of Teff production(quintal) was 24.23 quintal per household and the minimum and maximum Teff production(quintal) was 5 and 60 respectively, The previous studies reported that the average Teff production in Ethiopia was 8.78 quintal/hectare (24), Amhara regional state was 1.261 ton/hectare (25), 1.2 ton/hectare (26), 1.69 ton/hectare, Oromia region 1.717 ton/hectare, south nation nationality people region 1.38 and Benishanguel-Gumuz 1.24 ton/hectare (27). The observed differences in the productivity of Teff, in this study and other study conducted in different area could be different technology adoption, temperature (28), rainfall (29), climate (30), soil quality (31) and amount of nitrogen fertilizers used in growing the crop. The amount of DAP (kilogram) used in the previous year average of 72.44 kg/hectare. It was higher compared to the studies conducted in Ethiopia 29.1kg/hectare(24), Amhara regional state 66.36 kg/hectare (31). The mean figure of amount of UREA (kilogram/hectare) used was 47.07 kg/hectare (Table 1). This result lower than to the previous studies in Ethiopia 71.6 kg/hectare(24), Amhara regional state 52.85 kg/hectare (31). On the other hand the current result was higher than study in Ethiopia 38 kg/hectare(32). This is not consistent with the extension recommendations that require proper combination of DAP and UREA. Some works stated 100 kg of each of DAP and UREA per ha of cultivated land as a recommended (24).

With regard to gender of household heads, female headed households accounted for approximately 27.4% in both Teff technology adoption and non-adoption group and similarly male headed households were 72.6% (Table 2). The percentage indicates that the total respondents of male headed households were highest. Contributions of gender respondents were analyzed with respect to Teff technology adopter and non-adopter. Out of 72.6% (n=138) of the respondents male household head 81.2% were agricultural technology adopt and out of 27.4% (n=52) of the respondents female household head 69.2% agricultural technology adopt. This result implies that male households head participate more in the agricultural technology adoption activity than female household head. This difference assumed to be due to the labor intensive nature and high economy of agricultural technology adoption practices since women have extra responsibility in the house such as ensuring food security in their family, taking care of children and looking after livestock limits their participation in agricultural technology adoption practices. Gender of the household was a statistically significant with 95% confidence. This result consistence with the previous studies, for instance results of studies in sub-Saharan Africa have shown that male headed households have more access to land, education, and information on new technologies (33). In some countries female headed households are discriminated against by credit institutions, and as such they are unable to finance yield-raising technologies, leading to low adoption rates (34).

**Table 2:**
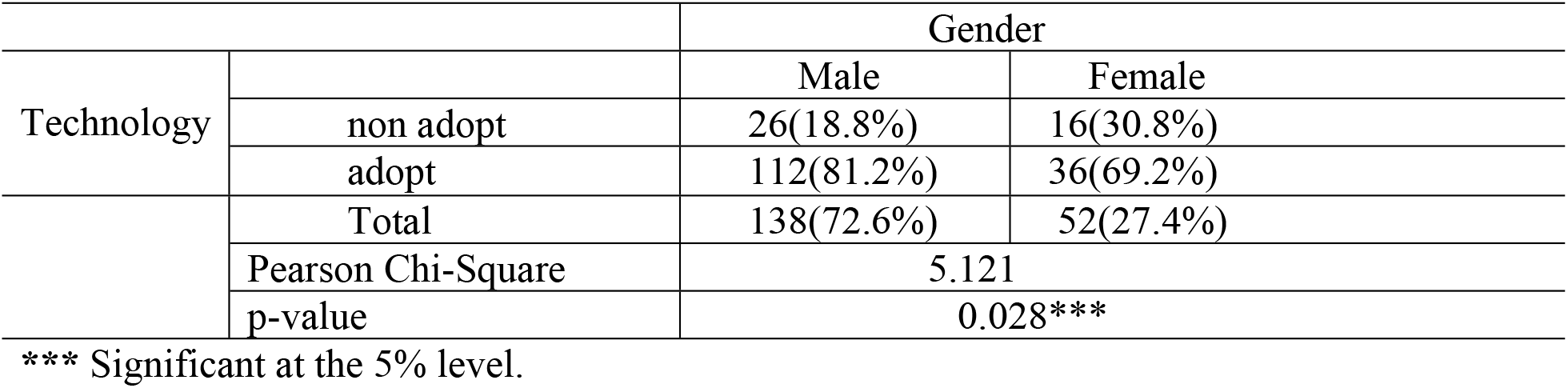
Gender composition of household heads and agricultural technology adoption

The educational level of household head differs with respect to the households’ in agricultural technology adopter, which may have a pronounced effect on Teff production. In the present study from the total respondent household head, 50% were illiterate with no formal educational background and the remaining 50% of the household head at least can read and write. From the total percentage of people who can read and write, the highest percent were primary education levels, i.e., 40%. From the total agricultural technology adopter households, 48.6%, 38.7%, 7.9% and 2.7% were illiterate, primary education level, secondary education level and college and above education level respectively. On the other hand, 54.8%, 45.2%, 0.0% and 0.00% of non-agricultural technology adopter households had education level of illiterate, primary school, secondary school and college and above respectively (Table 3). The non agriculture technology adopter group was more density in illiterate. This finding is consistence to the other study(9, 35). Education level of the household was an important characteristic in the study area. This is due to the fact education improves the access to information, new ideas and inputs provided by extension workers. Education may make a farmer more receptive to advice from an extension agency or more able to deal with technical recommendations that require a certain level of numeracy or literacy.

**Table 3:** Education level of the household head and agricultural technology adoption

| Variables | Category | Technology |  |  | Chi2-value | P –value |
| --- | --- | --- | --- | --- | --- | --- |
|  |  | non adopt | Adopt | Total |  |  |
| Education level | Illiterate | 23(54.8%) | 72(48.6%) | 95(50%) | 6.006 | 0.018*** |
|  | Primary | 19(45.2%) | 57(38.7%) | 76(40%) |  |  |
|  | Secondary | 0(0.0%) | 15(7.9%) | 15(7.9%) |  |  |
|  | College and above | 0(0.0%) | 4(2.7%) | 4(2.1%) |  |  |

\*\*\* Significant at the 5% level.

The average land size holding of agricultural technology adopter and non adopter household was 1.88 and 1.63 hectare respectively. The difference the average plot holding in agricultural technology adopter and non adopter is not large enough. The t-test indicates that the total land size is not different in agricultural technology adopter and non adopter group (Table 4). But by this minimum difference of land size or farm size the farmers have got highly different amount of products with the same land size. Even if the average farm size the same in agricultural technology adopter and non adopter, agricultural technology adopter households got high amount of Teff yield as compared to non agricultural technology adopter households.

**Table 4:** average landholding and Teff cultivated land

| Variables | Technology adoption |  | non technology adoption |  | T-value | P-value |
| --- | --- | --- | --- | --- | --- | --- |
|  | Mean | Std. deviation | Mean | Std. deviation |  |  |
| Total land size | 1.88 | 0.75 | 1.63 | 0.97 | -1.979 | 0.233*** |
| Land size under Teff cultivation | 1.83 | 1.10 | 1.60 | 0.54 | -1.195 | 0.971*** |

\*\*\* Significant at the 5% level.

Extension service means that giving training, advice, demonstration and distribution of agricultural inputs in to the agricultural farmers. The study shows that 87.9% of households get extension service. Contrary the survey result indicates that 12.1% from the total sampled households who was not got extension services. According to the survey study 100% technology adopter and 45.2% non-technology adopter households were get advices and supports from different agricultural agents. The result indicates that agricultural technology adopter household has got more extension services than non-adopter farmers. When we compare the access of extension services in agricultural technology adopter and non-adopter households’ who participate in Teff production activity, agricultural technology adopter has got more advices and support from the development agency or extension agents than to non-agricultural technology adopter. This result shows that agricultural technology adopter households got more extension service than non-adopter households. Agricultural technology practice requires close follow up to aware farmers in order to us chemical fertilizer, organic fertilizer, crop protected chemical and row planting. This result shows that the extension service is significant at 95% confidence interval among agricultural technology adopter and non-adopter (Table 5). This finding is agree with the studies conducted in Pakistan and Uganda (36, 37).

**Table 5:** Extension service and agricultural technology adoption

| extension service | Technology |  |  |  | Total |  | Chi2-value | p-value |
| --- | --- | --- | --- | --- | --- | --- | --- | --- |
|  | Adopter |  | Non- adopter |  |  |  |  |  |
| Yes | Freq. | % | Freq. | % | Freq. | % | 92.21 | 0.00*** |
|  | 148 | 100 | 19 | 45.2 | 167 | 87.9 |  |  |
| No | 0 | 0 | 23 | 54.8 | 23 | 12.1 |  |  |

\*\*\* Significant at the 5% level

Access to credit for farmers is important in order to purchase agricultural inputs like improved seed, DAP, UREA, chemicals for improving farm land, livestock’s. In the present study the proportion of the households who gets credit access was 34.2%. However, 65.8% of the sampled households did not participate in the credit service. According to this study 31.1% agricultural technology adopter and 45.2% non-agricultural technology adopter households were get credit access from different credit agents. This result shows that the credit access has significant effect on uses of agricultural technology at 95% confidence interval among agricultural technology adopter and non-adopter as chi-square=5.91 and p-value=0.03 indicates (Table 6).

**Table 6:** Credit Access and agricultural technology adoption

| Credit | Technology |  |  |  | Total |  | Chi2-value | p-value |
| --- | --- | --- | --- | --- | --- | --- | --- | --- |
|  | Adopter |  | Non- adopter |  |  |  |  |  |
| Yes | Freq. | % | Freq. | % | Freq. | % | 5.91 | 0.03*** |
|  | 46 | 31.1 | 19 | 45.2 | 65 | 34.2 |  |  |
| No | 102 | 68.9 | 23 | 54.8 | 125 | 65.8 |  |  |

\*\*\* Significant at the 5% level.

It is found that total land holding has a positive relastioship with teff production i.e r=0.47.The relatioship of total landholding of household and teff productivity indicates that the two variable has direct relashionship but it is wear relationship. When landholding incearses teff productivity also increase. The association is a linear increasing function form (supplementary Figure 1).

It is found that annual on farm income of the household head has a strong positive direct linear relastioship with teff production i.e r=0.99.The relatioship of annual on farm income of the household head and teff productivity indicates that the two variable has direct relashionship. When annual income of the household head incearses teff productivity also increse. The association is a linear increasing function form (supplemetary Figure 2).

The result of t-test clearly shows that there was no difference average Teff production in quintal between male household head and female household head with mean of 24.26 and 24.15 respectively. The amount of Teff produce by the household usually measure in quintal, one quintal means 100 kg. The result of t-test shows that there is no a significant difference Teff production in quintal between male and female household headed. P-value 0.96 shows that average Teff production has no significant difference based on sex in 95% confidence level (Table 7). This finding is inconsistent to a study done by Bisanda and Mwangi, (33) disclosed that male household head produce better than females and the results were statistically significant.

**Table 7:** The relation between Teff production and independent variables

|  | Variable |  | Mean mark | Std deviation | t | df | P value |
| --- | --- | --- | --- | --- | --- | --- | --- |
| Teff production | sex | male | 24.26 | 14.97 | 0.04 | 188 | 0.96 |
|  |  | female | 24.15 | 9.44 |  |  |  |
|  | Technology adoption | Non adopter | 14.85 | 5.55 | -5.40 | 188 | 0.00*** |
|  |  | adopter | 26.89 | 14.97 |  |  |  |
|  | Row planting | yes | 25.18 | 11.29 | -11.35 | 188 | 0.01*** |
|  |  | No | 21.94 | 16.78 |  |  |  |
|  | Credit | yes | 26.18 | 12.79 | -2.78 | 188 | 0.00*** |
|  |  | no | 20.47 | 13.73 |  |  |  |
|  | Extension service | yes | 25.48 | 13.95 | 3.50 | 188 | 0.00*** |
|  |  | no | 15.13 | 5.99 |  |  |  |
|  | Chemical fertilizer | yes | 25.48 | 13.95 | 3.50 | 188 | 0.00*** |
|  |  | no | 15.13 | 5.99 |  |  |  |
|  | Organic fertilizer | yes | 26.87 | 8.90 | -2.57 | 188 | 0.01*** |
|  |  | No | 21.85 | 17.16 |  |  |  |

\*\*\* Significant at the 5% level.

The average Teff productivity for household agriculture technology adopter was 26.89 quintal per hectare and for non-adopter was 14.85 quintal per hectare. This suggests that the productivity of Teff was greater in households adopting agricultural technology than in non-farm technology adopters. The T-value is -5.40 and the probability value is zero. In general the result tells that when household are more participate in agricultural technology adoption they become more technical efficient and productive. This could be attributed to various reasons, as they see the significance and implication of their knowledge with the help of extension service, they can apply the knowledge they have learned to use new context. Similarly, both technology adoption and households may be encouraged and accomplished by such a kind of particular education experience. After discussing the area of difficulty with the former to adopt technology, the extension service giver can offer an opportunity for continuous feedback. Indeed, extension service plays a vital role in improving households’ technology adoption, last Teff productivity becomes improve.

Average Teff production in quintal has a link with accessibility of credit for the household in the study area. After analyzing the result by using independent T test, the result shows that there is a significance difference between household who have no access to credit service and access to credit service on average Teff production in quintal. The mean Teff production of household with access to credit was 26.18 quintal/hectare and with non-access to credit have mean value of 20.47 quintal/hectare by support work gives T-value 4.22 and p-value is 0.00 (Table 7), this shows that household access credit service has a high Teff productivity than who does not access credit service.

Education level categorize as illiterate, primary, secondary and college and above. In this study the highest education level was college and above. The result of ANOVA one-way the P-value is 0.005.This result shows that there is significance difference between the Education level household head on average Teff productions in quintal per hectare (Table 8). To show which categories of education level average Teff production different from the other, the researcher conducted multiple comparisons using LSD. There is significance mean difference between no educated and secondary school and also secondary school with primary school but the rest pair of education level has a significance mean difference (supplementary Table 1).

**Table 8:** ANOVA for Teff productivity based on education level of household head

|  | Sum of square | Df | Mean square | F | sig |
| --- | --- | --- | --- | --- | --- |
| Between group | 2314.10 | 3 | 771.36 | 4.35 | 0.005 |
| Within group | 32961.71 | 186 | 177.21 |  |  |
| total | 35275.81 | 189 |  |  |  |

### Factors affecting households’ agricultural technology adoption

The outcome for household head gender, after adjusting for other covariates, indicates that the male household head was 7,644 times more likely than the reference group (female) household head in the study area to adopt agricultural technology and its effect has statistical significant (Table 9). These differences can be explained in part by limited access to productive resources due to tradition, culture and other institutional and economic constraints. This study conform the finding of different studies (25, 38-43).

**Table 9:** Final Logistic Regression of adoption of agricultural technology

| Covariate | Categories | $\hat{\beta}$ | S.E($\hat{\beta}$) | Wald | d.f. | P-value | Exp($\hat{\beta}$) |
| --- | --- | --- | --- | --- | --- | --- | --- |
| gender | male | 2.034 | 0.645 | .941 | 1 | 0.002*** | 7.644 |
|  | female | - | - | - | - | - | - |
| Family size |  | 0.139 | 0.284 | 2.556 | 1 | 0.025*** | 1.149 |
| age |  | -0.136 | 0.055 | 6.134 | 1 | 0.013*** | 0.873 |
| Row planting | yes | 5.550 | 0.917 | 36.598 | 1 | 0.000*** | 257.20 |
|  | no | - | - | - | - | - | - |
| Credit | yes | 1.145 | .869 | 2.734 | 1 | 0.001*** | 3.141 |
|  | no | - | - | - | - | - | - |
| Manure use | yes | -3.177 | 0.891 | 12.711 | 1 | 0.000*** | 0.042 |
|  | no | - | - | - | - | - | - |
| Constant |  | 1.658 | 1.865 | 0.790 | 1 | 0.374 | 5.247 |

\*\* \* significant at the 5% level.

As one unit rises in the age of the household head the chances of implementing agricultural technology were reducing by 0.873, keeping stable the other variables. This result is unexpected because, from empirical analysis, the age of the household head is usually taken as a proxy for farming experience. This finding confirm to study conducted in Ethiopia, Basso Liben district (9), this study showed that age of the household head had negative and significant effect on the adoption of Teff row planting practice at 1% significance level and the other study showed that on the average, older farmers are more likely to stop adopting the technology as their physical ability to participate actively in farming activities declines with increasing age(19, 31, 44-46). Older farmers may be less inclined to try new farm practices; younger farmers exposed to improved agricultural technologies will have increasing likelihood of agricultural technology adoption, ceteris paribus, as they become more aware of the benefits of agricultural technology adoption and have the opportunity to adjust productive resources over time.

Family size of the household has a positive and significance effect on the adoption of agricultural technology. The value of the coefficient *B* for a variable support is 0.139 with odd of 1.149; it entails that if one unit increases in family size, the odds of being adopting agricultural technology increase by 1.149 holding the other variables at constant. It is often assumed that farmers with larger family size will be more likely to adopt a new agricultural technology, especially if the innovation requires an extra cash investment. Familysize is also related to access to information or credit that would facilitate the agricultural technology adoption. Large family size may be an indicator for accessibility of labor provided that there are more people within the age range of active labor force. Therefore family size is expected to increase the probability of Teff technology adoption and Teff productivity in particular area. Household labor had positive and significant effect on the adoption of Teff row planting method at 5% significance level. The coefficient of farm labor represented by household size is statistically significant and has a positive association with technology adoption (9, 19, 20, 44, 47).

Credit access has direct relation with the adopting agricultural technology in the study area. A household head who have credit access were 3.141 times more likely to adopt agricultural technology than that of household head who have no credit access in Basso Liben district. Therefore the household head who have credit access more involve to adopt agricultural technology than the household head who have not credit access. If a recommendation implies significant cash investment for farmers, its adoption may be facilitated by an efficient credit program. If the majority of adopters use credit to acquire the agricultural technology, this is a strong indication of credit’s role in diffusing the technology. This finding is inconsistence with the study conducted in Ethiopia (9, 19, 20, 31).

Manure use in agriculture has indirect relation with the adopting agricultural technology. A household head that use manure in their Teff production were 0.042 times less likely to adopt agricultural technology than the household who do not use manure in their Teff production. The use of manure and compost has strong negative impact on the adoption of inorganic fertilizer (31).

### Factors affecting Teff productivity

Landholding has a positive and significance effect on the production of Teff (Table 10). This implies an increase in Teff productivity by 5.10 quintals if landholding increases by one unit. This finding confirm with the finding of Hailu et al. (48) on productivity of Teff varieties in Ethiopia.

**Table 10:** multiple regression model factors that affect Teff production

| Covariate | Categories | $\hat{\beta}$ | S.E($\hat{\beta}$) | t | p>\|t\| |
| --- | --- | --- | --- | --- | --- |
| age | age | 0.0791533 | 0.210 | 3.71 | 0.00*** |
| Education level | Illiterate | - | - | - | - |
|  | primary | -0.612759 | 0.246 | -2.48 | 0.014*** |
|  | secondary | -0.713204 | 0.445 | -1.60 | 0.111 |
|  | College and above | 0.0185444 | 0.803 | 0.02 | 0.982 |
| Total land |  | 5.1071082 | 0.174 | -4.63 | 0.000*** |
| Annual on income |  | 0.0005071 | 0.000 | 71.43 | 0.000*** |
| Off-farm income |  | 0.0000247 | 0.000 | 0.09 | 0.296 |
| Extension service | no | -0.635360 | 0.372 | -1.71 | 0.009*** |
|  | yes | - | - | - | - |
| Row planting | no | - | - | - | - |
|  | yes | 1.409936 | 0.255 | 5.51 | 0.000*** |
| Organic fertilizer | no | -0.946 | 0.281 | -3.36 | 0.001*** |
|  | yes | - | - | - | - |
| Constant |  | -2.124171 | 0.831 | -2.55 | 0.011*** |

\*\* \* significant at the 5% level.

The productivity of Teff was positively affected by age in this multiple regression model. It has been indicated that Teff production varies by vary the age of household head. This means that as age of household head increase by one unit, Teff productivity increase by 0.079 quintals. This is also statistically significance with p value 0.000. age of the household head is usually taken as a proxy for experience with farming this finding contradict to the study conducted in Ethiopia (48).

Annual income of the household head has a positively affected the productivity of Teff. It has been indicate that Teff production varies by vary the annual income of the household head. This means that if the annual income of household head increase by one unit Teff production increase by 0.0005 quintals held the other variables constant. This is statistically significant with p value 0.000. The income obtained from farm activities helps farmers to purchase farm inputs. Assessment of some of the previous empirical studies indicated that, the influence of annual income on Teff production varies from one finding to the other. But, majority of the studies reported positive contribution of annual income to household’s adoption of improved agricultural technologies.

Row planting has a positively affected the productivity of Teff. It has been indicate that Teff production varies by row planting user and non users. This means that if household head use row planting Teff production increase by 1.409936 quintals held the other variables constant. This is statistically significant with p value 0.000.

Extension service has a positively affected the productivity of Teff. It has been indicate that Teff productivity varies by extension service user and non users. This means that if household head not use extension service Teff productivity decrease by 0.63536 quintals held the other variables constant. This is statistically significant with p value 0.009.

Organic fertilizer has a positively affected the productivity of Teff. It has been indicate that Teff productivity varies by organic fertilizer user and non users. This means that if household head not use organic fertilizer productivity decrease by 0.946 quintals held the other variables constant. This is statistically significant with p value 0.001.

## CONCLUSIONS

The findings from this study show a low prevalence of agricultural technology adoption and Teff productivity in the study area and different associated agricultural technology adoption and Teff productivity factors have been identified. Thus, interventions by the bodies concerned on agricultural technology adoption and Teff productivity should ruminate extend credit access and agricultural extension service to cope with less agricultural technology adoption and Teff productive and also community-based education to access information on agricultural technology adoption and Teff productivity as well as high birth rates, which can increase family size. As manure, organic fertilizer and landholding are significant associated factors of agricultural technology adoption and Teff productivity, attention should be given to increase manure use, organic fertilizer use and mixed farming on the small available landholding to increase agricultural technology adoption and Teff productivity. Besides, row planting should be accomplished through extension service and awareness of row planting programs through farmer field schools. Finally, further study should be conducted to identify mechanisms for addressing agricultural technology adoption and Teff productivity in the study area.

## Acknowledgments

We thank Basso Liben district household for providing useful data. We also thank reviewers of this journal for their constructive comments.

## Author Contributions

Demelash A.: Designed the study, analyzed data and wrote the article and critically edited the manuscript.

Hayimero E.: Designed the study, analyzed data and edited the manuscript. All authors read and approved the final manuscript.

Mulusew K.: Designed the study, collect data, and edited the manuscript. All authors read and approved the final manuscript.

